# Susceptibility of domestic swine to experimental infection with SARS-CoV-2

**DOI:** 10.1101/2020.09.10.288548

**Authors:** Brad S. Pickering, Greg Smith, Mathieu M. Pinette, Carissa Embury-Hyatt, Estella Moffat, Peter Marszal, Charles E. Lewis

## Abstract

SARS-CoV-2, the agent responsible for COVID-19 has been shown to infect a number of species. The role of domestic livestock and the risk associated for humans in close contact remains unknown for many production animals. Determination of the susceptibility of pigs to SARS-CoV-2 is critical towards a One Health approach to manage the potential risk of zoonotic transmission. Here, pigs undergoing experimental inoculation are susceptible to SARS-CoV-2 at low levels. Viral RNA was detected in group oral fluids and nasal wash from at least two animals while live virus was isolated from a pig. Further, antibodies could be detected in two animals at 11 and 13 days post infection, while oral fluid samples at 6 days post inoculation indicated the presence of secreted antibodies. These data highlight the need for additional livestock assessment to better determine the potential role domestic animals may contribute towards the SARS-CoV-2 pandemic.

## Introduction

Severe acute respiratory syndrome coronavirus 2 (SARS-CoV-2), the agent of coronavirus disease (COVID-19) was recently identified to cause severe respiratory distress in humans with symptoms ranging from asymptomatic, mild to severe, and sometimes fatal cases (1). Rapidly spreading, this novel virus emerged in Wuhan China, to generate a pandemic as declared by the World Health Organization on March, 11^th^ 2020 (2). Predicted to have originated in bats, SARS-CoV-2 origins are still under intense investigation as reports continue to identify the ability of the virus to infect new animal species (3-8). Detection of natural infections has recently shed light on knowledge gaps in understanding transmission which has raised concerns regarding amplifying or reservoir hosts. In turn, a better understanding of wildlife and domestic animal susceptibility is required to assess the potential roles and present risks to prevent future spread of disease. Domestic swine, one of the most significant and highly produced agricultural species with previous impacts to public health, must be assessed (9-12). The increase in “backyard” small stakeholder animal production in both rural and urban environments provides an important source of high-quality protein and income, but can also serve as a source for zoonotic disease; therefore, it is important to investigate their potential role during SARS-CoV-2 spread (13). Evidence for the involvement of production animals was recently highlighted in The Netherlands where anthroponotic transmission of SARS-CoV-2 from humans to farmed mink with subsequent zoonotic transmission to at least two humans from mink has been proposed, further exemplifying the need to identify the potential role of production animals in disease transmission (14).

Angiotensin-converting enzyme 2 (ACE2) has been identified to be the receptor for SARS-CoV-2 (15). A Basic Local Alignment Search Tool (BLAST) query of the protein database using translated nucleotide (BLASTx) from the human ACE2 coding sequence predicts 98% coverage and 81% identity for the homologous receptor in swine. Interestingly, using the same search both mink (82%) and feline (85%) show similar identity to the human ACE2 for their cognate receptors. Moreover, both mink and cats have been reported to be susceptible to SARS-CoV-2 and have shown transmission to other animals (5, 16). Work by Zhou *et al*. utilized *in vitro* infectivity studies testing ACE2 receptor from laboratory mice, horseshoe bats, civets and the domestic pig. All of the respective receptors, except mice, were reported to enter HeLa cells indicating a functional target for SARS-CoV-2. Moreover, the authors employed additional known coronavirus receptors including both aminopeptidase N and dipeptidyl peptidase 4 finding neither are used for cell entry outlining the specificity for the ACE2 receptor (17).

The work reported here aims to determine whether domestic swine are susceptible to SARS-CoV-2 infection, providing critical information to aid public health risk assessments. Following oronasal inoculation, swine were assessed for: clinical signs and pathology, evidence of virus shedding, viral dissemination within tissues, and seroconversion. The data presented in this study provides evidence live SARS-CoV-2 virus can persist in swine for at least 13 days following experimental inoculation.

## Methods

### Ethics Statement

Experimental design, including housing conditions, sampling regimen, and humane endpoints, were approved by the Animal Care Committee of the Canadian Science Centre for Human and Animal Health in AUD #C-20-005 and all procedures and housing conditions were in strict accordance with the Canadian Council on Animal Care (CCAC) guidelines. Group housing was carried out in the BSL-3zoonotic large animal cubicles, and animals were provided with commercial toys for enrichment and access to food and water ad libitum. All invasive procedures, including experimental inoculation and sample collection (nasal washes, rectal swabs, and blood collection) were performed under isoflurane gas anesthesia, and animals were euthanized by intravenous administration of a commercial sodium pentobarbital solution.

### Study design

Nineteen domestic, American Yorkshire crossbred pigs (Sus scrufa) (6 castrated males and 13 females, age 8 weeks) were locally sourced in Manitoba, Canada and utilized in this study. Sixteen pigs were oronasally challenged with 1 × 10^6^ pfu per animal in a total of 3 mL DMEM under sedation with isoflurane. 1 ml was distributed per nostril and 1ml placed in the distal pharynx utilizing a sterile, tomcat-style catheter. The challenge dose was confirmed by back-titration of the inoculum on Vero E6 cells. Two naïve pigs were placed in the room with the inoculated pigs at day 10 to serve as in-room transmission controls. One additional uninoculated pig was sampled and necropsied to serve as a “farm control” providing negative control tissues. A physical examination including collection of blood, multiple swabs (rectal, oral, and nasal), and nasal wash with sterile D-PBS was performed at day zero and every other day beginning at three days post-inoculation (DPI) until day 15. The remaining pigs were sampled on 22 and 29 DPI. Group oral fluids from rope chews were collected daily. Necropsies and post-mortem sampling were performed starting at 3 DPI as outlined in Table 1. Animal numbers were not based on power analysis but on the limitations of the containment animal room size and requirements of CCAC guidelines. Group assignment (day of euthanasia and necropsy) was based on randomization at the time of permanent animal identification (ear tag).

**Table 1:** Sampling and necropsy schedule.

|  | Pig | 0 DPI | 3 DPI | 5 DPI | 7 DPI | 9 DPI | 11 DPI | 13 DPI | 15 DPI | 22 DPI | 29 DPI |
| --- | --- | --- | --- | --- | --- | --- | --- | --- | --- | --- | --- |
| cubicle 1 | 20-01 | + | + | - | - | - | - | - | - | - | - |
|  | 20-02 | + | + | + | - | - | - | - | - | - | - |
|  | 20-03 | + | + | + | + | - | - | - | - | - | - |
|  | 20-04 | + | + | + | + | + | - | - | - | - | - |
|  | 20-05 | + | + | + | + | + | + | - | - | - | - |
|  | 20-06 | + | + | + | + | + | + | + | - | - | - |
|  | 20-07 | + | + | + | + | + | + | + | + | - | - |
|  | 20-08 | + | + | + | + | + | + | + | + | + | - |
| cubicle 2 | 20-09 | + | + | - | - | - | - | - | - | - | - |
|  | 20-10 | + | + | + | - | - | - | - | - | - | - |
|  | 20-11 | + | + | + | + | - | - | - | - | - | - |
|  | 20-12 | + | + | + | + | + | - | - | - | - | - |
|  | 20-13 | + | + | + | + | + | + | - | - | - | - |
|  | 20-14 | + | + | + | + | + | + | + | - | - | - |
|  | 20-15 | + | + | + | + | + | + | + | + | - | - |
|  | 20-16 | + | + | + | + | + | + | + | + | + | + |
| contact animals | 20-17 | + | + | + | + | + | + | + | + | + | + |
|  | 20-18 | + | + | + | + | + | + | + | + | + | - |
|  | 20-19 | + | + | + | + | + | - | - | - | - | - |

\* indicates necropsy

### Sampling of animals

Oral, rectal, and nasal swabs were taken from each pig under general anesthesia using isoflurane and placed into sterile D-PBS containing the following antibiotics: streptomycin, vancomycin, nystatin, and gentamycin. Fluid was collected from a bilateral nasal wash using sterile D-PBS. Blood was collected in each of the following via jugular venipuncture: serum, sodium citrate, sodium heparin, and K3 EDTA.

Hematology, chemistry, and blood gas analyses. Hematology was performed on an HM5 analyzer (Abaxis) using K3 EDTA-treated whole blood and the following parameters were evaluated: red blood cells, hemoglobin, hematocrit, mean corpuscular volume, mean corpuscular hemoglobin, mean corpuscular hemoglobin concentration, red cell distribution weight, platelets, mean platelet volume, white blood cells, neutrophil count (absolute (abs) and %), lymphocyte count (abs and %), monocyte count (abs and %), eosinophil count (abs and %), and basophil count (abs and %). Blood chemistries were evaluated on a VetScan 2 (Abaxis) with the Comprehensive Diagnostic Profile rotor (Abaxis) using serum stored at −80°C until tested and the following parameters were evaluated: glucose, blood urea nitrogen, creatinine, calcium, albumin, total protein, alanine aminotransferase, aspartate aminotransferase, alkaline phosphatase, amylase, potassium, sodium, phosphate, chloride, globulin, and total bilirubin. Sodium heparin treated blood was used to analyze venous blood gases, which were performed on an iSTAT Alinity V machine (Abaxis) using a CG4+ cartridge (Abaxis) to measure the following parameters: lactate, pH, total carbon dioxide, partial pressure carbon dioxide, partial pressure oxygen, soluble oxygen, bicarbonate, and base excess. Age-specific values were utilized to establish normal ranges along with the machine reference intervals (18-20).

### Necropsy

Necropsies were performed after euthanasia via sodium pentobarbital overdose, confirmation of death, and exsanguination by femoral artery laceration. Tissue collection included the following, individually and split between 10% neutral-buffered formalin and fresh tissue: skeletal muscle, abdominal fat, liver, spleen, pancreas, duodenum, jejunum, ileum, spiral colon, kidney, gastrohepatic and mesenteric lymph nodes, right cranial lung lobe, right middle lung lobe, right caudal lung lobe, left cranial lung lobe, left caudal lung lobe, trachea, heart, tracheobronchial lymph nodes, cervical spinal cord, meninges, cerebrum, cerebellum, brainstem, olfactory bulb, nasal turbinates, submandibular lymph nodes, tonsil, trigeminal ganglion, and the entire eye. The reproductive tract (uterus and ovaries) were collected en bloc in female animals. Epiglottis and laryngeal folds were collected from some animals. The following were also collected at necropsy: cerebrospinal fluid, urine (when possible), vitreous, and bronchoalveolar lavage (BAL) utilizing Dulbecco’s Modified Eagle’s Medium (DMEM).

### Histopathology

Tissues were fixed in 10% neutral phosphate buffered formalin, routinely processed, sectioned at 5 µm and stained with hematoxylin and eosin (HE) for histopathologic examination.

### In situ hybridization

5 um paraffin-embedded formalin fixed tissue sections were ran according to the user manual for the RNAscope® 2.5HD Detection Reagent – Red kit by Advanced Cell Diagnostics (ACD) using the V-nCoV2019-S probe from ACD. The sections were then counter stained with Gill’s 1 hematoxylin, dried and coverslipped.

### Cells and virus

Second passage of SARS-CoV-2 (generously provided by the Public Health Agency of Canada, hCoV-19/Canada/ON-VIDO-01/2020) was propagated on Vero E6 cells in DMEM supplemented with 1% FBS. Virus was titrated by plaque assay and viral isolation was performed as previously described (21, 22).

Tissue homogenization and virus isolation. Pre-weighed, frozen tissue sections in Precellys bead mill tubes were thawed and D-PBS was added to make 10% w/v tissue homogenates. Tubes were processed using a Bertin Minilys personal tissue homogenizer and clarified by centrifugation at 2000 xg. Clarified homogenates, swabs and fluids collected from experimental animals were inactivated with TriPure Reagent (Roche) and extracted in duplicate as described below. Semi-quantitative real-time RT-PCR (RT-qPCR) positive samples were tested for virus isolation through standard plaque assay on VE6 cells using freshly prepared homogenates of frozen tissue.

### RNA Extraction

Total RNA from cell culture or experimental samples was extracted using the MagMax CORE Nucleic Acid Purification Kit per manufacturer’s recommendation with the following modifications. Briefly; sample was diluted in TriPure Reagent at a 1:9 ratio and used in place of the manufacturer’s lysis buffer for inactivation. 650 µL of TriPure-inactivated sample, 30µL of binding beads, and 350 µL of kit-provided CORE binding buffer spiked with Enteroviral armoured RNA was utilized followed by single washes in both Wash 1 and Wash 2 buffers, with a final elution volume of 30 μL of kit-supplied elution buffer using the automated MagMax Express 96 system running the KingFisher-96 Heated Script “MaxMAX_CORE_KF-96” (ThermoFisher Scientific, 2020). The spiked Enteroviral armoured RNA (ARM-ENTERO; Asuragen) was used as an exogenous extraction and reaction control.

### Detection of SARS-CoV-2

RT-qPCR was performed on all extracted samples including sodium citrate whole blood using SARS-CoV-2 Envelope (E) gene specific primers and probe (23). RT-qPCR was utilized to detect viral RNA using the following primers and probe: E_SARBECO_F1 FOR: 5’-ACAGGTACGTTAATAGTTAATAGCGT-3’, E_SARBECO_R2 REV: 5’-ATATTGCAGCAGTACGCACACA-3’, and E_SARBECO-P1 Probe: 5’-ACACTAGCCATCCTTACTGCGCTTCG-3’. Mastermix for RT-qPCR was prepared using 4X TaqMan® Fast Virus 1-step master mix according to manufacturer’s specifications, using 0.4 µM of each E gene primer and 0.2 µM of probe per reaction. Reaction conditions were as follows: 50°C for 5 min, 95°C for 20 sec, and 40 cycles of 95°C for 3s followed by 60°C for 30s. Runs were performed using a 7500 Fast Real-Time PCR System (Thermofisher, ABI) and semi-quantitative results were calculated based on a gBlock (Integrated DNA Technologies, IDT) standard curve for SARS-CoV-2 E gene. SARS-CoV-2 specific primers were used for confirmation targeting both the Spike gene (S gene); SARS2_Spike_Probe: 5’TGCCACCTTTGCTCACAGATGAAATGA-3’, SARS2_Spike_FOR:5’-TGATTGCCTTGGTGATATTGCT-3’, SARS2_Spike_REV: 5’CGCTAACAGTGCAGAAGTGTATTGA-3’ and the RNA dependent RNA Polymerase gene (RdRp gene) RdRp_SARSr-F 5’-GTGARATGGTCATGTGTGGCGG-3’ RdRp_SARSr-R 5’CARATGTTAAASACACTATTAGCATA-3’ and RdRp_SARSr-P2 5’-CAGGTGGAACCTCATCAGGAGATGC-3’.

### Genome sequencing

Extracted RNA from the Submandibular lymph node of pig 20-06 was processed for high-throughput sequencing with enrichment for sequences for vertebrate viruses according to previously published method and sequenced on an Illumina MiSeq Reagent Kit v3 (600-cycle) (1, 24). Sequences were analysed using an in-house nf-villumina (v2.0.0) Nextflow workflow that performed: read quality filtering with fastp; taxonomic classification with Centrifuge and Kraken2 using a Centrifuge index of NCBI nt downloaded 2020-02-04 and a Kraken2 index of NCBI RefSeq sequences of archaea, bacteria, viral and the human genome GRCh38 downloaded and built on 2019-03-22; removal of non-viral reads (NCBI taxonomic id 10239) using the Kraken2 and Centrifuge taxonomic classification results; de novo assembly by Shovill, Unicycler and Megahit using the taxonomically filtered reads; nucleotide BLAST search of all assembled contigs against NCBI nucleotide downloaded 2020-04-10. nf-villumina taxonomically filtered reads were mapped against the top viral nucleotide BLAST match (SARS-CoV-2 isolate 2019-nCoV/USA-CA3/2020, MT027062.1) to generate a consensus sequence.

### Serum Neutralization Assays

Neutralizing antibody titers in sera were determined via plaque reduction neutralization test against SARS-CoV-2. Serial five-fold dilutions of heat inactivated (30 min at 56°C) sera were incubated with virus for 1 hour at 37°C. Each virus-serum mixture was then added to duplicate wells of Vero E6 cells in a 48-well format, incubated for 1 hour at 37°C, and overlaid with 500 μl of 2.0% carboxymethylcellulose in DMEM per well. Plates were then incubated at 37°C for 72 hours, fixed with 10% buffered formalin, and stained with 0.5% crystal violet. Serum dilutions resulting in >70% reduction of plaque counts compared to virus controls were considered positive for virus neutralization.

### Surrogate virus Neutralization Test

Detection and semi-quantitation of neutralizing antibodies was determined using SARS-CoV-2 Surrogate Virus Neutralization Test Kit (Genscript, Cat. No.: L00847). Testing of sera was performed as outlined in the manufacturer’s instructions. All sera were assessed from 7 DPI through 29 DPI including archived negative sera and kit-supplied negative controls. Neutralization was determined positive when above the recommended 20% cutoff.

## Results

Experimental inoculation of sixteen eight week old swine was performed oronasally with 1 × 10^6^ pfu of SARS-CoV-2, distributed evenly between both nostrils and the distal pharynx. Starting at 1 day post inoculation (DPI), pigs developed a mild, bilateral ocular discharge and in some cases, this was accompanied by serous nasal secretion. This was observed for only the first three days post inoculation. Temperatures remained normal throughout the study (Table S1). Overall, animals did not develop clinically observable respiratory distress, however one animal (Pig 20-06) presented mild depression at 1 DPI accompanied with a cough which was maintained through 4 DPI. This animal did not display additional clinical signs over the course of the study.

Viral shedding was evaluated to identify potential transmission which may occur through droplets from coughing, sneezing, oral fluids or gastrointestinal involvement. A sampling schedule was developed with the goal of determining the incidence of viral shedding (Table 1). Every other day starting at 3 DPI to 15 DPI, oral, nasal, and rectal swabs were sampled to evaluate the potential for delayed onset (1). Nucleic acid was extracted from swabs and RT-qPCR was performed to identify SARS-CoV-2 by targeting the envelope gene (E gene). Viral RNA could not be detected in swabs from any animals over the course of the study (Table 2, A). Nasal washes are a sensitive method for detection of pathogens in swine and were routinely sampled using sterile D-PBS to rinse nasal passages. Two pigs (20-10, 20-11) displayed low levels of viral RNA by RT-qPCR at 3 DPI (Table 2, A). Recovery of live virus was attempted for both PCR-positive nasal wash samples, however neither produced cytopathic effect or increased RNA detection via RT-qPCR of the cell culture supernatent. A third method for the detection of viral shedding was performed using a non-invasive, group sampling method. A cotton rope was hung on the pen prior to feeding which allows for deposit of oral fluids. Daily fluids from ropes were processed, with one room (Table 1, Cubicle 1) a weak positive signal for viral RNA at 3 DPI by RT-qPCR. Virus isolation was attempted from this sample, but similar to nasal washes, virus was not successfully isolated. It is important to note the positive oral fluid from cubicle 1 is not from the same room as the two positive nasal washes generated from Pigs 20-10 and 20-11 as the later were in different animal cubicles. Therefore, at minimum three animals provide evidence of viral nucleic acid in oronasal secretions from two independent animal rooms. Of note, two naïve pigs were introduced to the infected pigs at 10 DPI as transmission contacts, however no indication of viral infection could be detected from these animals at any point during the study.

**Table 2:** Detection of SARS-CoV-2 by real-time RT-PCR.

**A**
|  | 0 DPI | 3 DPI | 5 DPI | 5 DPI | 9 DPI | 11 DPI | 13 DPI | 15 DPI | 22 DPI | 29 DPI |
| --- | --- | --- | --- | --- | --- | --- | --- | --- | --- | --- |
| Swabs | 0\57 | 0\57 | 0\51 | 0\45 | 0\39 | 0\30 | 0\24 | 0\18 | 0\12 | 0\6 |
| Nasal Wash | 0\16 | 2\16 | 0\14 | 0\12 | 0\12 | 0\10 | 0\8 | 0\6 | 0\4 | 0\2 |
| Blood | 0\16 | 0\16 | 0\14 | 0\12 | 0\12 | 0\10 | 0\8 | 0\6 | 0\4 | 0\2 |

**B**
|  | 0 DPI | 1 DPI | 2 DPI | 3 DPI | 4 DPI | 5 DPI | 6 DPI | 7 DPI | 8 DPI | 9 DPI | 10 DPI | 11 DPI | 12 DPI | 13 DPI | 14 DPI |
| --- | --- | --- | --- | --- | --- | --- | --- | --- | --- | --- | --- | --- | --- | --- | --- |
| Oral fluids | 0\2 | 0\2 | 0\2 | 1\2 | 0\2 | 0\2 | 0\2 | 0\2 | 0\2 | 0\2 | 0\2 | 0\2 | 0\2 | 0\2 | 0\2 |
|  | 15 DPI | 16 DPI | 17 DPI | 18 DPI | 19 DPI | 20 DPI | 21 DPI | 22 DPI | 23 DPI | 24 DPI | 25 DPI | 26 DPI | 27 DPI | 28 DPI | 29 DPI |
| Oral fluids | 0\2 | 0\2 | 0\2 | 0\2 | 0\2 | 0\2 | 0\2 | 0\1 | 0\1 | 0\1 | 0\1 | 0\1 | 0\1 | 0\1 | 0\1 |

**C**
|  | 3 DPI |  | 5 DPI |  | 7 DPI |  | 9 DPI |  | 11 DPI |  | 13 DPI |  | 15 DPI |  | 22 DPI |  | 29 DPI |  |
| --- | --- | --- | --- | --- | --- | --- | --- | --- | --- | --- | --- | --- | --- | --- | --- | --- | --- | --- |
| Pig ID | 20-01 | 20-09 | 20-02 | 20-10 | 20-03 | 20-11 | 20-04 | 20-12 | 20-05 | 20-13 | 20-06 | 20-14 | 20-07 | 20-15 | 20-08 | 20-18 | 20-16 | 20-17 |
| no. tissues | 0\35 | 0\35 | 0\36 | 0\35 | 0\35 | 0\35 | 0\35 | 0\34 | 0\35 | 0\36 | 1\35 | 0\35 | 0\36 | 0\35 | 0\36 | 0\35 | 0\35 | 0\35 |

\* grey cells indicate positive RT-qPCR

Detection of SARS-CoV-2 was also attempted from whole blood by RT-qPCR, following the sampling schedule outlined in Table 1. As outlined in Table 2A, viremia, as indicated by the presence of viral RNA in the blood, could not be detected in any animal throughout the study. Blood cell counts, chemistries, and gasses were measured using the Abaxis HM5, VetScan 2, and iSTAT respectively. Although some variation was observed throughout the study, changes were minimal and inconclusive, and profiles consistent with acute viral infection or subsequent organ damage were not observed.

To identify potential target tissues or gross lesions consistent with SARS-CoV-2 disease, necropsy was performed on two animals starting at 3 DPI and every other day up to day 15; with an additional two pigs necropsied at both 22 and 29 DPI (Table 1). No significant pathology was observed which could be directly attributed to a viral infection. RT-qPCR was performed across all tissues and samples collected at necropsy targeting the E gene of SARS-CoV-2 (Table 2, C). One tissue, the submandibular lymph node tested from Pig 20-06 necropsied at 13 DPI was positive for viral RNA. The tissue sample was repeated in triplicate, on independent days, generating consistent results. Further, RNA was extracted from homogenized tissue and full genome sequence of SARS-CoV-2 was recovered. A 10% homogenate was generated from the submandibular lymph node and used to infect Vero E6 cells. Aliquots were taken from cell culture at 2DPI and 3DPI to monitor viral replication as indicated by an increasing quantity of RNA. Mild CPE was observed by 3DPI in the first passage with an increase in viral RNA measured by RT-qPCR targeting the envelope, spike and RNA dependent RNA polymerase genes. The first passage supernatant was clarified by centrifugation and passaged a second time in Vero E6 cells. At 2DPI of the second passage, significant CPE was exhibited in addition to increasing copies of SARS-CoV-2 viral RNA confirmed by RT-qPCR. Together this demonstrated the presence of live, replication-competent SARS-CoV-2 virus isolated form the submandibular lymph node of Pig 20-06 (Table 2).

The development of SARS-CoV-2 neutralizing antibodies were monitored over the course of study. Starting at 7 DPI, serum was obtained from individual animals for both virus neutralization test (traditional VNT) and a surrogate virus neutralization test (sVNT; Genscript). Sera was first tested using a traditional VNT, with one pig (20-07) generating neutralizing antibody titers, albeit weak, at a 1:5 dilution with a 70% reduction of plaques at both 13 and 15 DPI (Table 3). Consequently, the sVNT assay identified the same animal, Pig 20-07, as antibody positive with 0.188 µg/ml antibody at 15 DPI. A second pig (20-14) was shown to have generated antibody at 11 DPI (0.113 µg/ml) and 13 DPI (0.224 µg/ml). The sVNT was also employed to identify secreted antibody in oral fluids throughout the study. Interestingly, at 6 DPI we detected positive antibody (0.133 µg/ml) from group oral fluid collected from cubicle 1 (Table 3).

**Table 3:** Neutralizing antibody development.

| Pig | 0 DPI |  | 7 DPI |  | 9 DPI |  | 11 DPI |  | 13 DPI |  | 15 DPI |  | 22 DPI |  | 29 DPI |  |
| --- | --- | --- | --- | --- | --- | --- | --- | --- | --- | --- | --- | --- | --- | --- | --- | --- |
|  | VNT<br>(1:5) | sVNT<br>(µg/ml) | VNT<br>(1:5) | sVNT<br>(µg/ml) | VNT<br>(1:5) | sVNT<br>(µg/ml) | VNT<br>(1:5) | sVNT<br>(µg/ml) | VNT<br>(1:5) | sVNT<br>(µg/ml) | VNT<br>(1:5) | sVNT<br>(µg/ml) | VNT<br>(1:5) | sVNT<br>(µg/ml) | VNT<br>(1:5) | sVNT<br>(µg/ml) |
| 20-01 | - | - | - | - | - | - | - | - | - | - | - | - | - | - | - | - |
| 20-02 | - | - | - | - | - | - | - | - | - | - | - | - | - | - | - | - |
| 20-03 | - | - | - | - | - | - | - | - | - | - | - | - | - | - | - | - |
| 20-04 | - | - | - | - | - | - | - | - | - | - | - | - | - | - | - | - |
| 20-05 | - | - | - | - | - | - | - | - | - | - | - | - | - | - | - | - |
| 20-06 | - | - | - | - | - | - | - | - | - | - | - | - | - | - | - | - |
| 20-07 | - | - | - | - | - | - | - | - | + | - | + | 0.188 | - | - | - | - |
| 20-08 | - | - | - | - | - | - | - | - | - | - | - | - | - | - | - | - |
| 20-09 | - | - | - | - | - | - | - | - | - | - | - | - | - | - | - | - |
| 20-10 | - | - | - | - | - | - | - | - | - | - | - | - | - | - | - | - |
| 20-11 | - | - | - | - | - | - | - | - | - | - | - | - | - | - | - | - |
| 20-12 | - | - | - | - | - | - | - | - | - | - | - | - | - | - | - | - |
| 20-13 | - | - | - | - | - | - | - | - | - | - | - | - | - | - | - | - |
| 20-14 | - | - | - | - | - | - | - | 0.113 | - | 0.224 | - | - | - | - | - | - |
| 20-15 | - | - | - | - | - | - | - | - | - | - | - | - | - | - | - | - |
| 20-16 | - | - | - | - | - | - | - | - | - | - | - | - | - | - | - | - |
| 20-17 | - | - | - | - | - | - | - | - | - | - | - | - | - | - | - | - |
| 20-18 | - | - | - | - | - | - | - | - | - | - | - | - | - | - | - | - |
| 20-19 | - | - | - | - | - | - | - | - | - | - | - | - | - | - | - | - |
| sVNT (µg/ml) |  |  |  |  |  |  |  |  |  |  |  |  |  |  |  |  |
|  | 0 DPI | 1 DPI | 2 DPI | 3 DPI | 4 DPI | 5 DPI | 6 DPI | 7 DPI | 8 DPI | 9 DPI | 10 DPI | 11 DPI | 12 DPI | 13 DPI | 14 DPI | 15 DPI |
| Oral Fluid | - | - | - | - | - | - | 0.113 | - | - | - | - | - | - | - | - | - |

## Discussion

The results presented in this study define domestic swine as a susceptible species albeit at low levels to SARS-CoV-2 viral infection. One animal was found to retain live virus, while two additional animals had detectible RNA measured in the nasal wash, and two pigs developed antibodies. In total, of the sixteen animals experimentally inoculated, five displayed some level of exposure or elicited an immune response to the virus, representing roughly 30% of the study cohort. One pig displayed mild, non-specific clinical signs, including coughing and depression in addition to multiple pigs demonstrating mild ocular and nasal discharge. These signs occurred during what could be considered to be the immediate, post-infection period. Over a nine day period, between cessation of clinical signs and post mortem evaluation, the virus was found to be maintained undetected in the submandibular lymph node in this animal. Importantly, of the five animals indicated potential infection, viral RNA was only detected at low levels and no live viral shedding was identified.

Following the detection of viral RNA in group oral fluids collected by rope chews at 3 DPI, the presence of secreted antibody was detected using a surrogate virus neutralization test (sVNT) at 6 DPI in the same sample type. The amount of antibody measured in oral fluids from swine would be considered below a protective cutoff based on comparisons to classical neutralizing titers, however the discovery of secreted antibody in oral fluids may be a useful tool for surveillance efforts. This also demonstrates the possibility that human saliva should be evaluated as a less invasive method to provide accompanying evidence with serosurveillance studies for exposure to SARS-CoV-2.

The results of this study contradict previous reports indicating swine are not susceptible to SARS-CoV-2 infection (4, 25). RNA was not detected in swabs or organ samples and no seroconversion was measured in these studies. Infectious dose, viral isolate, age, and breed or colony of swine may affect study outcomes. It should be noted in this work, a ten-fold higher viral dose was utilized for experimental infection compared to previous studies. Moreover, animals were obtained from a high health status farm in Manitoba, in contrast to a specific pathogen free colony, for the purpose of determining the risk to Canadian pigs. Altogether, these findings indicate that further investigations into the susceptibility of additional domestic livestock species should be studied to assess their risk. Finally, we emphasize that no cases of domestic livestock have been documented by natural infection to date; however, the results of this study support further investigations in to the role that animals may play in the maintenance and spread of SARS-CoV-2.

## Acknowledgments

We would like to thank the Public Health Agency of Canada for SARS-CoV-2 isolate for this study, in addition the Animal Care and Genomics units for their support during this project. We would also like to thank Dr. Claire Andreasen for her review of the clinical pathology findings.

## Author contributions

B.P. conceived the research. B.P, G.S., M.M.P., E.M, P.M., and C.E.L. performed the experiments. B.P, G.S., M.M.P., C.E.H and C.E.L. analysed the data. B.P. wrote the manuscript, with input from B.P, G.S., M.M.P., E.M, and C.E.L. All authors discussed the results and reviewed the manuscript. Competing interests: Authors declare no competing interests. Data and materials availability: all data is available in the manuscript or the supplementary materials.

## Biographical Sketch

Brad Pickering is the Head of the Special Pathogens Unit at the National Centre for Foreign Animal Disease with the Canadian Food Inspection Agency. His research focuses on high consequence pathogens including both emerging and re-emerging zoonotic diseases of veterinary importance.

## References

1. Li Q, Guan X, Wu P, Wang X, Zhou L, Tong Y, et al. Early Transmission Dynamics in Wuhan, China, of Novel Coronavirus–Infected Pneumonia. 2020;382(13):1199–207.

2. WHO. “WHO Director-General’s opening remarks at the media briefing on COVID19 -March 2020.” Retrieved June 14, 2020, from https://www.who.int/dg/speeches/detail/who-director-general-s-opening-remarks-at-the-media-briefing-on-covid-1911-march-2020. 2020 March

3. Sia SF, Yan LM, Chin AWH, Fung K, Choy KT, Wong AYL, et al. Pathogenesis and transmission of SARS-CoV-2 in golden hamsters. Nature. 2020 May 14.

4. Shi J, Wen Z, Zhong G, Yang H, Wang C, Huang B, et al. Susceptibility of ferrets, cats, dogs, and other domesticated animals to SARS-coronavirus 2. Science (New York, NY). 2020 May 29;368(6494):1016–20.

5. Oreshkova N, Molenaar R-J, Vreman S, Harders F, Munnink BBO, Hakze R, et al. SARS-CoV2 infection in farmed mink, Netherlands, April 2020. bioRxiv. 2020:2020.05.18.101493.

6. Lam TT, Shum MH, Zhu HC, Tong YG, Ni XB, Liao YS, et al. Identifying SARS-CoV-2 related coronaviruses in Malayan pangolins. Nature. 2020 Mar 26.

7. Andersen KG, Rambaut A, Lipkin WI, Holmes EC, Garry RF. The proximal origin of SARS-CoV-2. Nature medicine. 2020 Apr;26(4):450–2.

8. Abdel-Moneim AS, Abdelwhab EM. Evidence for SARS-CoV-2 Infection of Animal Hosts. Pathogens. 2020 Jun 30;9(7).

9. Smith TC, Harper AL, Nair R, Wardyn SE, Hanson BM, Ferguson DD, et al. Emerging Swine Zoonoses. Vector-Borne and Zoonotic Diseases. 2011 2011/09/01;11(9):1225–34.

10. Chua KB, Bellini WJ, Rota PA, Harcourt BH, Tamin A, Lam SK, et al. Nipah virus: a recently emergent deadly paramyxovirus. Science (New York, NY). 2000 May 26;288(5470):1432–5.

11. Chua KB, Goh KJ, Wong KT, Kamarulzaman A, Tan PS, Ksiazek TG, et al. Fatal encephalitis due to Nipah virus among pig-farmers in Malaysia. Lancet (London, England). 1999 Oct 9;354(9186):1257–9.

12. Mansfield KL, Hernández-Triana LM, Banyard AC, Fooks AR, Johnson N. Japanese encephalitis virus infection, diagnosis and control in domestic animals. Veterinary Microbiology. 2017 2017/03/01/;201:85–92.

13. Roth JA. Veterinary Vaccines and Their Importance to Animal Health and Public Health. Procedia in Vaccinology. 2011 2011/01/01/;5:127–36.

14. Enserink M. Coronavirus rips through Dutch mink farms, triggering culls. Science (New York, NY). 2020;368(6496):1169.

15. Letko M, Marzi A, Munster V. Functional assessment of cell entry and receptor usage for SARS-CoV-2 and other lineage B betacoronaviruses. Nat Microbiol. 2020 Apr;5(4):562–9.

16. Halfmann PJ, Hatta M, Chiba S, Maemura T, Fan S, Takeda M, et al. Transmission of SARS-CoV-2 in Domestic Cats. N Engl J Med. 2020 May 13.

17. Zhou P, Yang XL, Wang XG, Hu B, Zhang L, Zhang W, et al. A pneumonia outbreak associated with a new coronavirus of probable bat origin. Nature. 2020 Mar;579(7798):270–3.

18. Friendship RM, Lumsden JH, McMillan I, Wilson MR. Hematology and biochemistry reference values for Ontario swine. Can J Comp Med. 1984;48(4):390–3.

19. Perri AM, O’Sullivan TL, Harding JCS, Wood RD, Friendship RM. Hematology and biochemistry reference intervals for Ontario commercial nursing pigs close to the time of weaning. Can Vet J. 2017;58(4):371–6.

20. Ventrella D, Dondi F, Barone F, Serafini F, Elmi A, Giunti M, et al. The biomedical piglet: establishing reference intervals for haematology and clinical chemistry parameters of two age groups with and without iron supplementation. BMC Vet Res. 2017;13(1):23-.

21. Weingartl HM, Berhane Y, Caswell JL, Loosmore S, Audonnet JC, Roth JA, et al. Recombinant nipah virus vaccines protect pigs against challenge. J Virol. 2006 Aug;80(16):7929–38.

22. Li M, Embury-Hyatt C, Weingartl HM. Experimental inoculation study indicates swine as a potential host for Hendra virus. Vet Res. 2010 May-Jun;41(3):33.

23. Corman VM, Landt O, Kaiser M, Molenkamp R, Meijer A, Chu DK, et al. Detection of 2019 novel coronavirus (2019-nCoV) by real-time RT-PCR. Euro Surveill. 2020 Jan;25(3).

24. Zhang YZ, Holmes EC. A Genomic Perspective on the Origin and Emergence of SARS-CoV-2. Cell. 2020 Apr 16;181(2):223–7.

25. Schlottau K, Rissmann M, Graaf A, Schön J, Sehl J, Wylezich C, et al. SARS-CoV-2 in fruit bats, ferrets, pigs, and chickens: an experimental transmission study. The Lancet Microbe. 2020 2020/07/07/.

