## Supplementary tables for "Susceptibility of domestic swine to experimental infection with SARS-CoV-2"

The tables below include physiological parameters during samples under general anesthetic including; temperature, pulse and blood oxygenation. Supplementary table 1 is referrenced in text, while supplementary tables 2 and 3 are not.

### Supplementary table 1: Pig daily rectal body temperature

| Pig ID | | -3 | -2 | -1 | 0* | 1 | 2 | 3 | 4 | 5 | 6 | 7 | 8 | 9 | 10 | 11 | 12 | 13 | 14 | 15 | 16 | 17 | 18 | 19 | 20 | 21 | 22 | 23 | 24 | 25 | 26 | 27 | 28 | 29 | DPI |
|---|---|---|---|---|---|---|---|---|---|---|---|---|---|---|---|---|---|---|---|---|---|---|---|---|---|---|---|---|---|---|---|---|---|---|---|
| Animal Cubicle 1 | 20-01 | 39.8 | 39.8 | 39.4 | 39.4 | 39.7 | 39.3 | 39.3 | | | | | | | | | | | | | | | | | | | | | | | | | | | Rectal Body Temperature (degree celsius) |
| | 20-02 | 39.9 | 39.6 | 39.4 | 39.1 | 39.3 | 39.6 | 38.8 | 40.1 | 39.1 | | | | | | | | | | | | | | | | | | | | | | | | | |
| | 20-03 | 39.8 | 39.3 | 39.3 | 38.7 | 39.7 | 39.0 | 38.5 | 39.3 | 38.7 | 38.9 | 38.5 | | | | | | | | | | | | | | | | | | | | | | | |
| | 20-04 | 39.4 | 39.8 | 39.1 | 39.3 | 39.7 | 39.3 | 38.8 | 39.4 | 38.9 | 39.4 | 38.9 | 39.4 | 38.8 | | | | | | | | | | | | | | | | | | | | | |
| | 20-05 | 39.8 | 39.3 | 39.7 | 39.2 | 39.9 | 39.7 | 39.1 | 39.4 | 39.4 | 39.4 | 38.8 | 39.8 | 39.2 | 40.0 | 39.1 | | | | | | | | | | | | | | | | | | | |
| | 20-06 | 39.6 | 39.3 | 39.3 | 39.2 | 39.3 | 39.1 | 39.0 | 39.3 | 39.6 | 39.1 | 39.2 | 39.6 | 39.4 | 39.7 | 39.3 | 39.4 | 38.6 | | | | | | | | | | | | | | | | | |
| | 20-07 | 39.1 | 39.1 | 39.2 | 39.0 | 39.8 | 39.5 | 38.8 | 39.4 | 39.3 | 39.5 | 38.9 | 39.6 | 39.1 | 40.1 | 38.8 | 39.1 | 38.8 | 39.6 | 38.8 | | | | | | | | | | | | | | | |
| | 20-08 | 40.1 | 40.1 | 39.7 | 39.6 | 39.9 | 39.5 | 38.7 | 40.0 | 39.7 | 39.7 | 39.1 | 39.8 | 38.8 | 40.2 | 39.2 | 39.6 | 39.2 | 39.7 | 39.8 | 40.2 | 39.6 | 40.2 | 39.9 | 40.2 | 40.0 | 40.0 | | | | | | | | |
| | 20-18 | 40.0 | 39.5 | 39.8 | 39.2 | 39.2 | 39.5 | 39.0 | 39.0 | 39.0 | 39.4 | 39.0 | 38.9 | 39.3 | 38.9 | 38.5 | 39.6 | 38.9 | 39.7 | 39.1 | 39.2 | 39.2 | 39.5 | 38.6 | 39.0 | 39.7 | 39.7 | | | | | | | | |
| Animal Cubicle 2 | 20-09 | 39.7 | 39.6 | 40.0 | 39.2 | 39.6 | 39.9 | 39.0 | | | | | | | | | | | | | | | | | | | | | | | | | | | |
| | 20-10 | 39.7 | 39.6 | 39.9 | 39.4 | 39.7 | 39.6 | 38.7 | 39.8 | 39.4 | | | | | | | | | | | | | | | | | | | | | | | | | |
| | 20-11 | 39.7 | 40.2 | 39.3 | 39.3 | 39.7 | 39.1 | 39.2 | 39.9 | 39.6 | 39.4 | 38.6 | | | | | | | | | | | | | | | | | | | | | | | |
| | 20-12 | 39.8 | 39.6 | 39.8 | 39.4 | 39.1 | 39.0 | 38.8 | 39.9 | 39.7 | 39.6 | 38.6 | 39.3 | 39.4 | | | | | | | | | | | | | | | | | | | | | |
| | 20-13 | 39.8 | 39.9 | 40.1 | 39.0 | 39.8 | 39.7 | 39.3 | 39.8 | 39.6 | 39.6 | 39.1 | 40.0 | 39.9 | 40.3 | 39.7 | | | | | | | | | | | | | | | | | | | |
| | 20-14 | 39.7 | 39.7 | 39.8 | 39.4 | 39.1 | 40.0 | 39.6 | 39.9 | 39.7 | 39.1 | 38.7 | 39.9 | 39.5 | 40.1 | 39.7 | 39.4 | 39.0 | | | | | | | | | | | | | | | | | |
| | 20-15 | 39.6 | 39.3 | 39.5 | 39.6 | 39.5 | 39.4 | 38.4 | 39.5 | 39.6 | 38.9 | 38.5 | 39.6 | 39.6 | 39.6 | 39.5 | 39.7 | 39.4 | 39.6 | 38.8 | | | | | | | | | | | | | | | |
| | 20-16 | 39.7 | 39.2 | 39.0 | 38.9 | 39.4 | 39.4 | 39.0 | 39.9 | 39.7 | 39.3 | 38.7 | 39.7 | 39.2 | 39.9 | 39.3 | 39.9 | 38.8 | 39.6 | 39.5 | 39.8 | 39.6 | 39.3 | 39.8 | 39.2 | 39.5 | 39.2 | 39.2 | 39.3 | 39.5 | 39.8 | 39.8 | 39.8 | 38.8 | |
| | 20-17 | 39.4 | 39.6 | 39.5 | 39.5 | 39.4 | 38.7 | 39.3 | 38.5 | 39.6 | 39.4 | 39.8 | 38.4 | 39.1 | 38.8 | 39.0 | 39.1 | 38.4 | 39.5 | 38.6 | 39.5 | 39.1 | 39.3 | 39.7 | 39.9 | 38.9 | 39.4 | 38.9 | 39.5 | 40.2 | 39.0 | 39.0 | 39.9 | 39.4 | |
| FC | 20-19 | 39.6 | 39.0 | 39.5 | 39.5 | 39.3 | 39.5 | 39.6 | 38.9 | 39.1 | 39.3 | 39.5 | 38.1 | 38.9 | 38.8 | | | | | | | | | | | | | | | | | | | | |

Farm Control, FC.
*0 DPI is the day of inoculation

### Supplementary table 2: Pig aural pulse during general anesthesia.

| Pig ID | | -3 | -2 | -1 | 0* | 1 | 2 | 3 | 4 | 5 | 6 | 7 | 8 | 9 | 10 | 11 | 12 | 13 | 14 | 15 | 16 | 17 | 18 | 19 | 20 | 21 | 22 | 23 | 24 | 25 | 26 | 27 | 28 | 29 | DPI |
|---|---|---|---|---|---|---|---|---|---|---|---|---|---|---|---|---|---|---|---|---|---|---|---|---|---|---|---|---|---|---|---|---|---|---|---|
| Animal Cubicle A | 20-01 | | | | 183 | | | 195 | | | | | | | | | | | | | | | | | | | | | | | | | | | Pulse (beats per minute) |
| | 20-02 | | | | 189 | | | 99 | | 162 | | | | | | | | | | | | | | | | | | | | | | | | | |
| | 20-03 | | | | 162 | | | 155 | | 160 | | 144 | | | | | | | | | | | | | | | | | | | | | | | |
| | 20-04 | | | | 214 | | | 184 | | 170 | | 170 | | 179 | | | | | | | | | | | | | | | | | | | | | |
| | 20-05 | | | | 179 | | | 195 | | 221 | | 173 | | 198 | | 196 | | | | | | | | | | | | | | | | | | | |
| | 20-06 | | | | 180 | | | 208 | | 213 | | 168 | | 198 | | 185 | | 172 | | | | | | | | | | | | | | | | | |
| | 20-07 | | | | 190 | | | 193 | | 197 | | 167 | | 202 | | 170 | | 170 | | 205 | | | | | | | | | | | | | | | |
| | 20-08 | | | | 221 | | | 232 | | 234 | | 234 | | 228 | | 234 | | 227 | | 221 | | | | | | | 237 | | | | | | | | |
| | 20-18 | | | | | | | | | | | | | | | 128 | | 145 | | 159 | | | | | | | 208 | | | | | | | | |
| Animal Cubicle B | 20-09 | | | | 193 | | | 208 | | | | | | | | | | | | | | | | | | | | | | | | | | | |
| | 20-10 | | | | 188 | | | 230 | | 221 | | | | | | | | | | | | | | | | | | | | | | | | | |
| | 20-11 | | | | 195 | | | 219 | | 222 | | 198 | | | | | | | | | | | | | | | | | | | | | | | |
| | 20-12 | | | | 198 | | | 206 | | 197 | | 199 | | 197 | | | | | | | | | | | | | | | | | | | | | |
| | 20-13 | | | | 195 | | | 173 | | 234 | | 237 | | 228 | | 181 | | | | | | | | | | | | | | | | | | | |
| | 20-14 | | | | 222 | | | 221 | | 234 | | 213 | | 219 | | 234 | | 235 | | | | | | | | | | | | | | | | | |
| | 20-15 | | | | 158 | | | 173 | | 132 | | 180 | | 179 | | 170 | | 190 | | 159 | | | | | | | | | | | | | | | |
| | 20-16 | | | | 220 | | | 208 | | 188 | | 202 | | 153 | | 188 | | 193 | | 183 | | | | | | | 250 | | | | | | | 208 | |
| | 20-17 | | | | | | | | | | | | | | | 221 | | 144 | | 199 | | | | | | | 240 | | | | | | | 220 | |
| FC | 20-19 | | | | | | | | | | | | | | | | | | | | | | | | | | | | | | | | | | |

Farm Control, FC.
*0 DPI is the day of inoculation

### Supplementary table 3: Pig blood oxygenation as measured by pulse oximetry under general anesthesia

| Pig ID | | -3 | -2 | -1 | 0* | 1 | 2 | 3 | 4 | 5 | 6 | 7 | 8 | 9 | 10 | 11 | 12 | 13 | 14 | 15 | 16 | 17 | 18 | 19 | 20 | 21 | 22 | 23 | 24 | 25 | 26 | 27 | 28 | 29 | DPI |
|---|---|---|---|---|---|---|---|---|---|---|---|---|---|---|---|---|---|---|---|---|---|---|---|---|---|---|---|---|---|---|---|---|---|---|---|
| Animal Cubicle A | 20-01 | | | | 97% | | | 100% | | | | | | | | | | | | | | | | | | | | | | | | | | | Percent oxygen saturation |
| | 20-02 | | | | 100% | | | 142% | | 99% | | | | | | | | | | | | | | | | | | | | | | | | | |
| | 20-03 | | | | 99% | | | 100% | | 100% | | 99% | | | | | | | | | | | | | | | | | | | | | | | |
| | 20-04 | | | | 100% | | | 100% | | 98% | | 100% | | 100% | | | | | | | | | | | | | | | | | | | | | |
| | 20-05 | | | | 100% | | | 99% | | 100% | | 100% | | 100% | | 100% | | | | | | | | | | | | | | | | | | | |
| | 20-06 | | | | 100% | | | 100% | | 100% | | 100% | | 100% | | 100% | | 98% | | | | | | | | | | | | | | | | | |
| | 20-07 | | | | 99% | | | 100% | | 100% | | 100% | | 99% | | 98% | | 100% | | 100% | | | | | | | | | | | | | | | |
| | 20-08 | | | | 99% | | | 100% | | 100% | | 100% | | 99% | | 100% | | 99% | | 100% | | | | | | | 98% | | | | | | | | |
| | 20-18 | | | | | | | | | | | | | | | 98% | | 100% | | 100% | | | | | | | 100% | | | | | | | | |
| Animal Cubicle B | 20-09 | | | | 100% | | | 100% | | | | | | | | | | | | | | | | | | | | | | | | | | | |
| | 20-10 | | | | 100% | | | 100% | | 100% | | | | | | | | | | | | | | | | | | | | | | | | | |
| | 20-11 | | | | 100% | | | 100% | | 100% | | 100% | | | | | | | | | | | | | | | | | | | | | | | |
| | 20-12 | | | | 100% | | | 100% | | 99% | | 100% | | 100% | | | | | | | | | | | | | | | | | | | | | |
| | 20-13 | | | | 100% | | | 100% | | 100% | | 99% | | 99% | | 99% | | | | | | | | | | | | | | | | | | | |
| | 20-14 | | | | 100% | | | 100% | | 100% | | 100% | | 98% | | 98% | | 100% | | | | | | | | | | | | | | | | | |
| | 20-15 | | | | 100% | | | 100% | | 96% | | 100% | | 99% | | 98% | | 100% | | 100% | | | | | | | | | | | | | | | |
| | 20-16 | | | | 98% | | | 100% | | 100% | | 96% | | 100% | | 100% | | 100% | | 100% | | | | | | | 100% | | | | | | | 98% | |
| | 20-17 | | | | | | | | | | | | | | | 96% | | 100% | | 100% | | | | | | | 97% | | | | | | | 99% | |
| FC | 20-19 | | | | | | | | | | | | | | | | | | | | | | | | | | | | | | | | | | |

Farm Control, FC.
*0 DPI is the day of inoculation
